## Supplemental Figures for "Casein Kinases 2-dependent phosphorylation of the placental ligand VAR2CSA regulates Plasmodium falciparum-infected erythrocytes cytoadhesion"

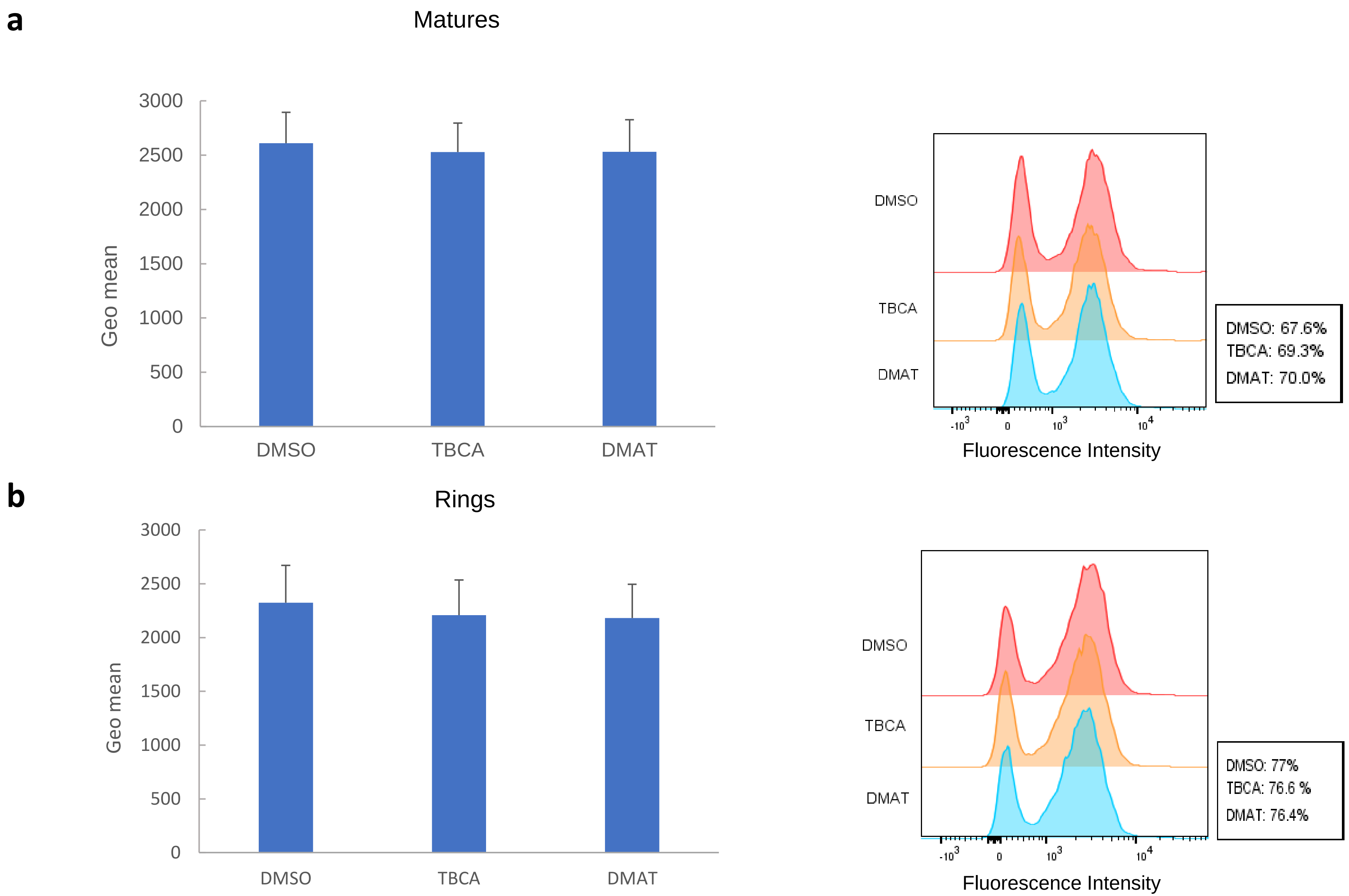


**Supplementary Figure 1. VAR2 surface expression and trafficking**

**(A)**. VAR2CSA Surface expression of trophozoite IEs after one treatment with CK2 inhibitors or DMSO monitored by flow cytometry with a specific anti-VAR2CSA antibody. **(B)** VAR2CSA Surface expression of ring stage IEs after 16 hours treatment with CK2 inhibitors or DMSO monitored by flow cytometry with a specific anti-VAR2CSA antibody geometric means of fluorescence intensities of three independent experiments are represented with standard deviations.

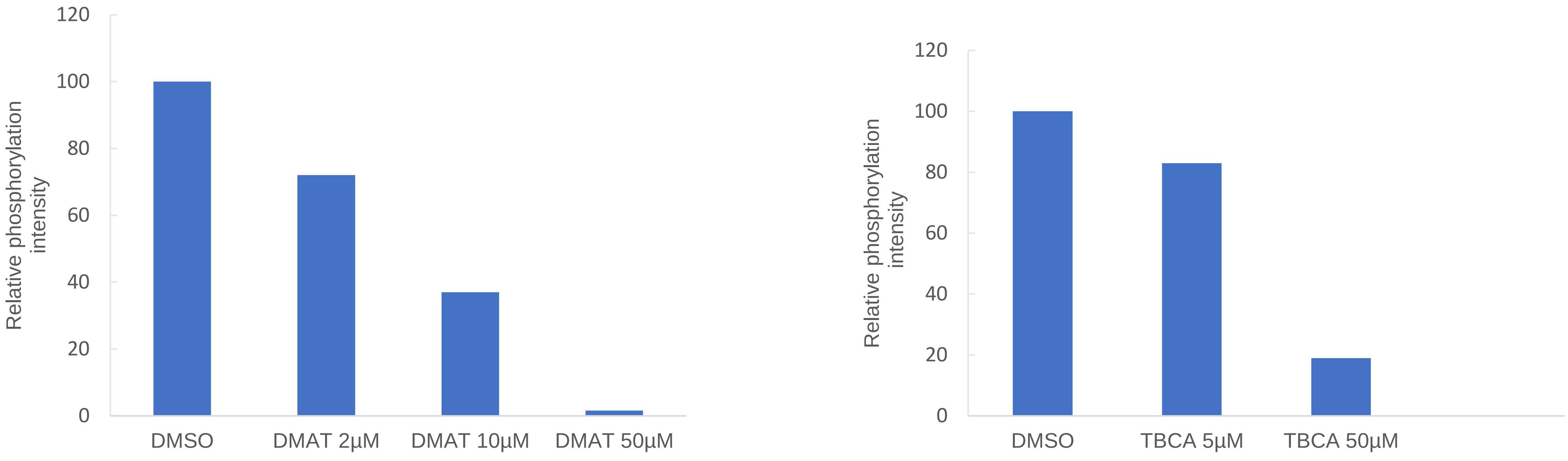


**Supplementary Figure 2. Quantification of phosphorylation signal**

The phosphorylation signal for each condition was quantified by Image Lab Software and adjusted to reflect a percentage compared to signal obtained in DMSO control condition (100%)

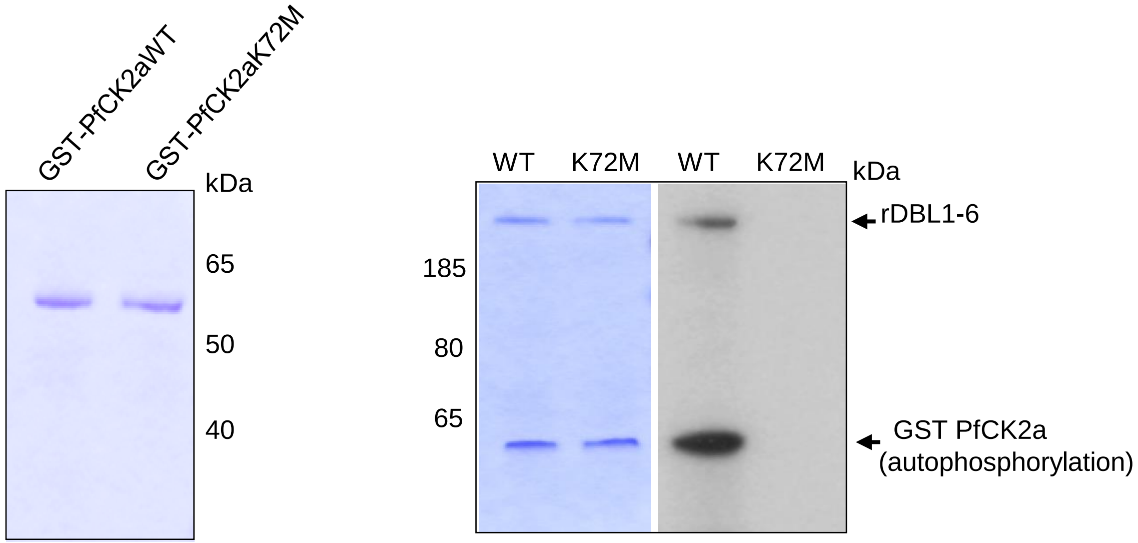


**Supplementary Figure 3. Expression and kinase activity of GSTPfCK2α and GSTPfCK2αK72M**

GSTPfCK2α and GSTPFCK2α K72M were expressed in *E. Coli* Rosetta and purified as described in [27] GST-PfCK2α kinase activity towards rDBL1-6. Autoradiograms (right) and Coomassie blue-stained gels (left) of kinase assays performed with GST-PfCK2α or catalytically inactive GST-K72MPfCK2α The recombinant kinase and rDBL1-6 substrate are indicated with an arrow. Autophosphorylation of the wild-type kinase is shown.

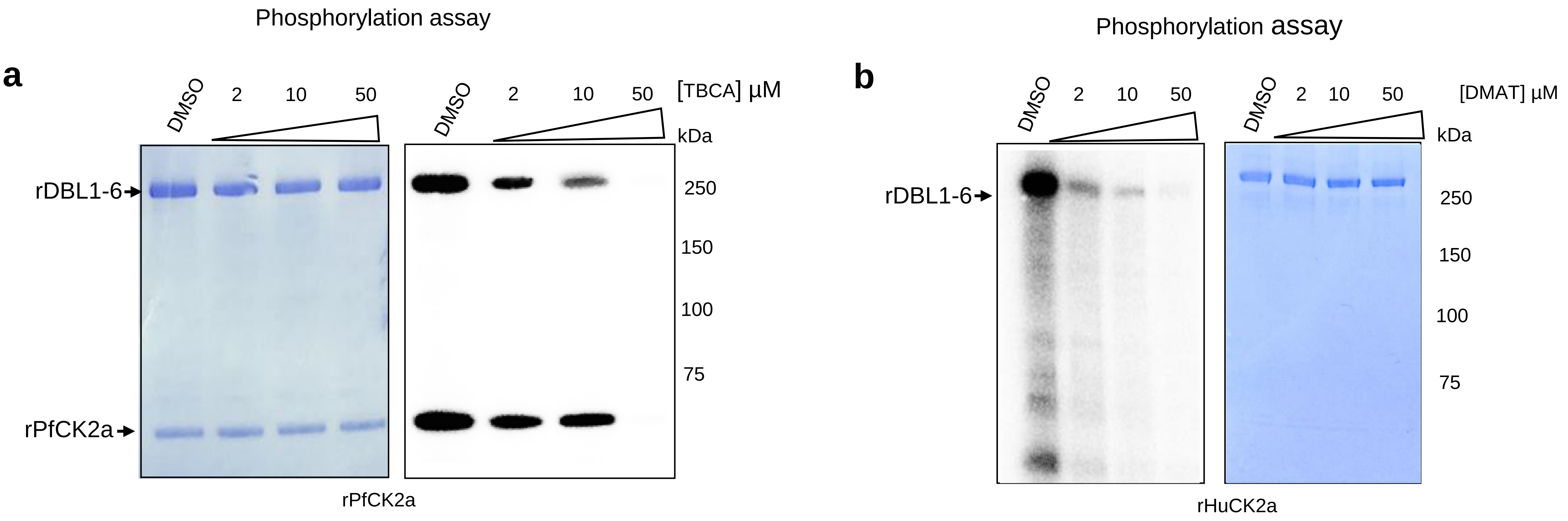


**Supplementary Figure 4. DMAT and TBCA dose dependent inhibition of *in vitro* rDBL1-6 phosphorylation by recombinant kinases**

Recombinant DBL1-6 was used in *in vitro* phosphorylation assays in the presence of [γ - 32P] ATP with recombinant Plasmodium kinase or Human CK2α and with increasing concentrations of DMAT and TBCA. **Panel A** (PfCK2α +TBCA): lane1: rDBL1-6 + DMSO; lane2: rDBL1-6 + TBCA 2µM; lane 3: rDBL1-6 + TBCA 10µM; lane 4: rDBL1-6 + TBCA 50µM; **Panel B** (HuCK2α + DMAT): lane1: rDBL1-6 + DMSO; lane2: rDBL1-6 + DMAT 2µM; lane 3: rDBL1-6 + DMAT 10µM; lane 4: rDBL1-6 + DMAT 50µM

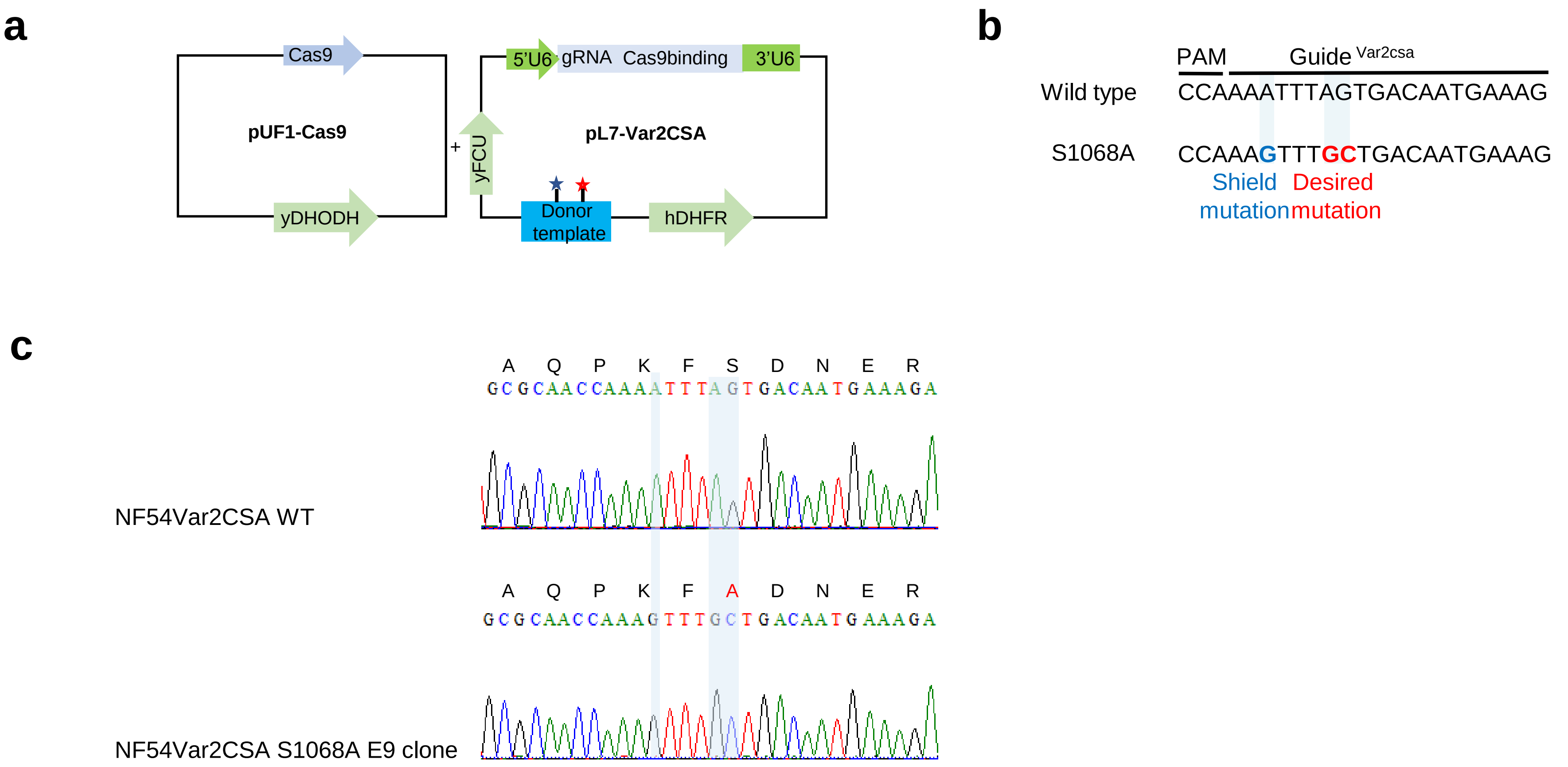


**Supplementary Figure 5. CRISPR/Cas9 strategy used for S1068 substitution.**

Nucleotide editing using sgRNA: Cas9 in *P. falciparum.* **A.** Diagram illustrating the strategy used for nucleotide replacement. The Cas9 protein is expressed in the pUF1-Cas9 episome continuously maintained using the ydhodh drug-selectable marker. PL7-VAR2CSA episome is maintained using the hdhfr selection and carries both the sgRNAVAR2CSA and the donor DNA (blue box). The donor DNA carries the designed desired mutation (red star) and the shield mutation (blue star). **B.** sgRNA VAR2CSA targeted sequences recognized by Cas9. The 20 nucleotides guide and PAM sequences are indicated. **C.** Chromatograms showing sequence analyses of VAR2CSA locus in NF54CSA wild-type and in transgenic NF54CSA S1068A E9 clone. Nucleotide substitutions and amino acids changes in VAR2CSA locus are highlighted

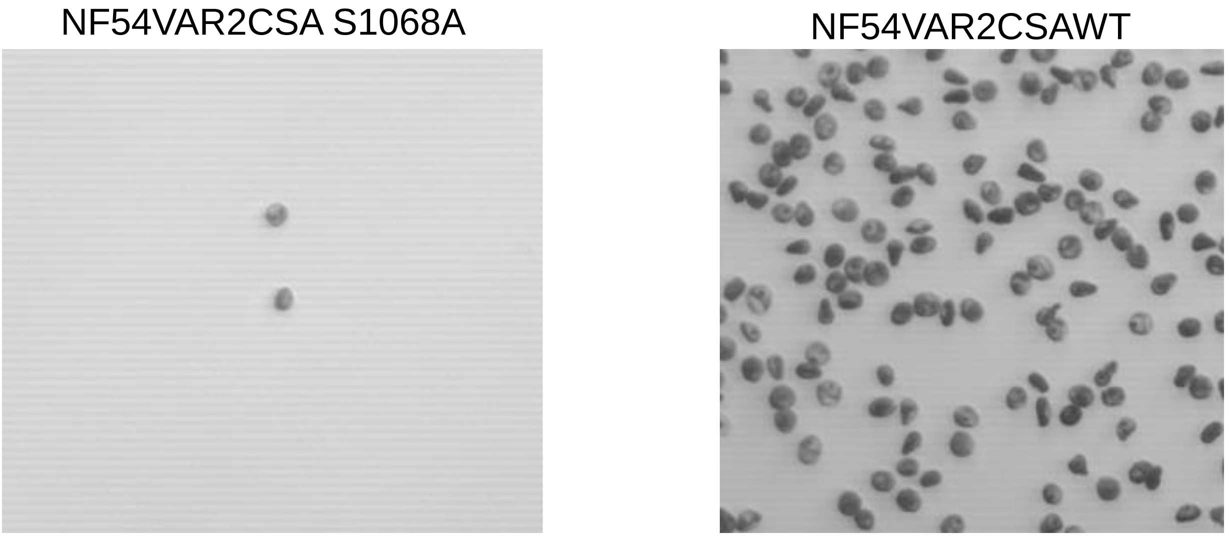


**Supplementary Figure 6. Static adhesion assay**. Image of a field showing bound NF54VAR2CSA parental and transgenic S1068A IEs on CSA coated on plastic.
